# Lipocalin-2 is a Sensitive and Specific Marker of Bacterial Infection in Children

**DOI:** 10.1101/623819

**Authors:** Jethro Herberg, Honglei Huang, Marie L. Thezenas, Victoria Janes, Michael Carter, Stuart Gormley, Melisa S. Hamilton, Benedikt Kessler, Michael Levin, Climent Casals-Pascual

## Abstract

**Introduction:** Bacterial infection is the leading cause of death in children globally. Clinical algorithms to identify children who are likely to benefit from antimicrobial treatment remain suboptimal. Biomarkers that accurately identify serious bacterial infection (SBI) could improve diagnosis and clinical management. Lipocalin 2 (LCN2) and neutrophil collagenase (MMP-8) are neutrophil-derived biomarkers associated with bacterial infection.

**Methods:** We evaluated LCN2 and MMP-8 as candidate biomarkers in 40 healthy controls and 151 febrile children categorised confirmed SBI, probable SBI, or viral infection. The diagnostic performance of LCN2 and MMP-8 to predict SBI was estimated by the area under the receiver operating characteristic curve (AUROC) and compared to the performance of C-reactive protein (CRP).

**Results:** Plasma LCN2 and MMP-8 concentration were predictive of SBI. The AUROC (95% CI) for LCN2, MMP8 and CRP to predict SBI was 0.88 (0.82-0.94); 0.80 (0.72-0.87) and 0.89 (0.84-0.94), respectively. The diagnostic performance of LCN2 in combination with CRP was significantly superior to either marker alone: AUROC 0.92 (95% CI: 0.88-0.96).

**Conclusion:** LCN2 is a sensitive and specific predictor of SBI in children which could be used to improve clinical management and antimicrobial stewardship. LCN2 should be further evaluated in prospective clinical studies.

## Introduction

Bacterial infection remains the major infectious cause of death in children (1). Identifying the small number of children with serious bacterial infection (SBI-microbiologically confirmed bacterial infection with systemic symptoms) amongst the majority with viral infections remains difficult, as clinical features are unreliable. Laboratory markers including white cell count, neutrophil proportion and C-reactive protein (CRP) have the strongest predictive value in children with clinically obvious SBI, rather than in the more numerous group with clinically uncertain infection status (2). Blood cultures, considered the gold standard for diagnosing SBI, are unhelpful for early antibiotic decisions as results are not immediately available, and are often negative if pre-admission antibiotics have been administered. In the UK there has been a 30% increase in hospital admission for children with probable infection (3), and the majority of children admitted with fever are discharged without a pathogen diagnosis. Many children with self-limiting viral illnesses are admitted to hospital and receive unnecessary antibiotics whilst awaiting the results of cultures, contributing to the spread of antibiotic resistance (4). There is an urgent need for new biomarkers that can reliably identify SBI.

We identified lipocalin-2 (LCN2, neutrophil gelatinase-associated lipocalin, NGAL) and neutrophil collagenase (matrix metalloproteinase-8, MMP-8) as potential protein candidates to distinguish bacterial pneumonia from viral pneumonia in African children (5). Here, we validate the diagnostic performance of these markers in a cohort of UK children with confirmed, probable or possible SBI versus viral infection and healthy controls.

## Methods

### Patient recruitment

Between July 2009 and April 2012, we recruited acutely ill febrile children (age <17 years) presenting with illness of sufficient severity to warrant blood tests. The study had approval of the St Mary’s Research Ethics Committee (REC 09/H0712/58). Written, informed consent was obtained.

### Pathogen diagnosis

Bacterial diagnostics included culture of blood and (when clinically indicated) cerebrospinal and pleural fluid, and pneumococcal antigen detection in blood or urine. Respiratory or nasopharyngeal secretions were screened for viruses by nested PCR, including RSV, coronavirus, adenovirus, parainfluenza 1-4, influenza A+B, bocavirus, metapneumovirus and rhinovirus.

### Patient categorisation and selection

Patients were assigned to clinical categories corresponding to likelihood of infection after evaluation of all clinical, radiological and laboratory data (Figure 1). Patient categories were ‘Definite Bacterial’ (DB), ‘Probable Bacterial’ (PB), ‘Uncertain Bacterial or Viral’ (U), ‘Probable Viral’ (PV) or ‘Definite Viral’ (DV). DB children had microbiologically confirmed bacterial infection identified at a sterile-site, irrespective of the CRP value. As there were only 4 patients in the PV category, analysis was not carried out on this group. Controls (C) were recruited in out-patients. After categorising, patients were prioritised for biomarker estimation based on the quantity of available plasma, and the chronological order of recruitment to give 5 groups as follows: 42 (DB), 34 (PB), 33 (U), 42 (V), 40 (C).

**Fig. 1.**
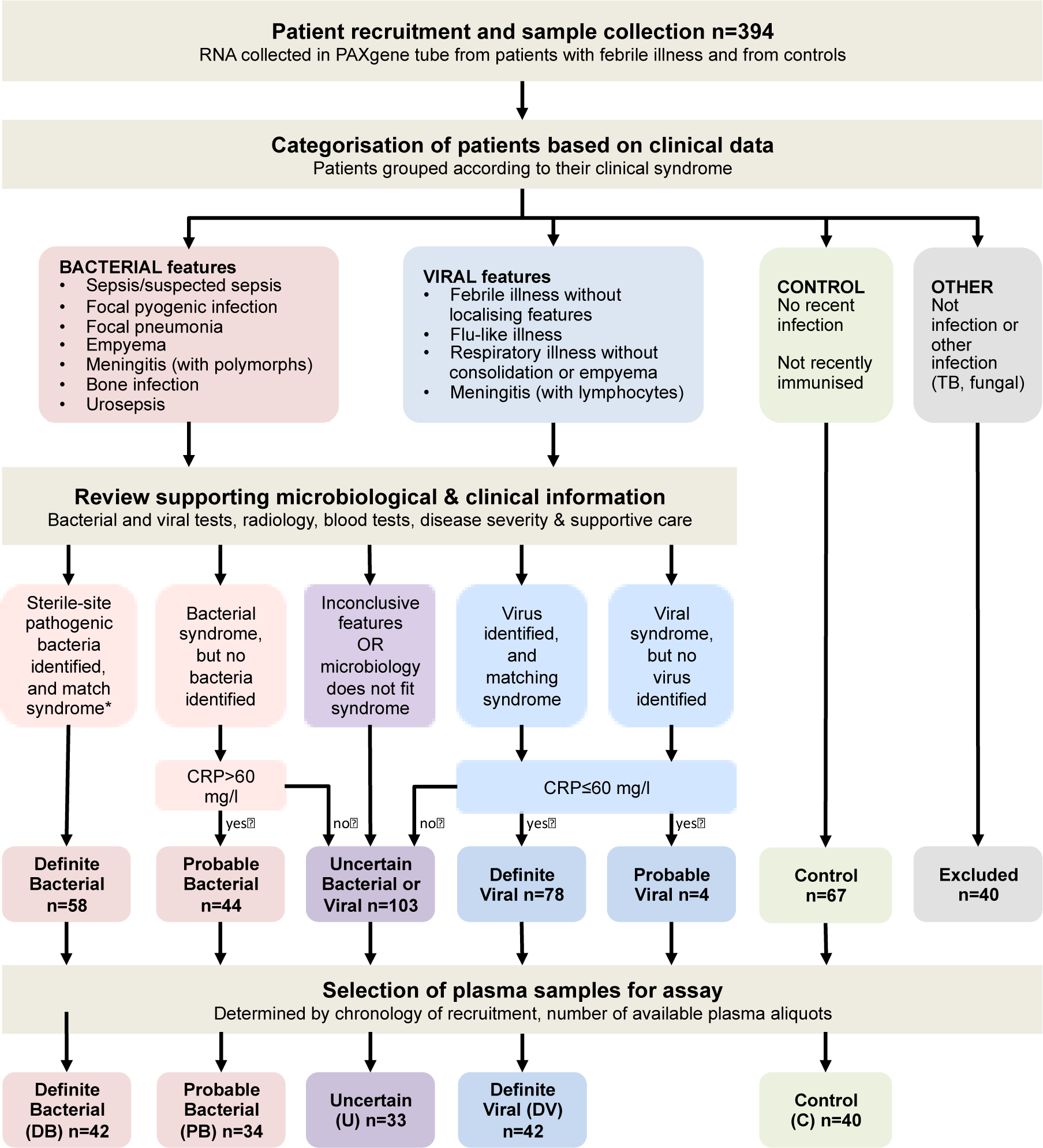
Clinical classification of the population studied: Assigning microbiological aetiology. Children admitted with acute infection, sick enough to warrant blood tests, were recruited after presentation and before diagnostic studies were completed, at St Mary’s Hospital, London and University Hospital NHS Foundation Trust, Southampton, UK. Detailed clinical and laboratory data were recorded. Healthy controls were recruited in out-patients. After excluding patients with proven or possible inflammatory conditions or mycobacterial disease, two independent paediatric infectious disease clinicians with access to all clinical and diagnostic data categorised patients as follows. Children with a clinical syndrome in keeping with SBI (sepsis with shock or severe focal infection) were categorised as ‘Definite Bacterial’ (DB) only if pathogenic bacteria were detected at a usually sterile site such as blood or CSF, and not including surface swabs, endotracheal secretions or bronchoalveolar lavage samples; otherwise these patients were categorised as ‘Probable Bacterial’ (PB). Children with a clinical syndrome in keeping with viral infection, not displaying any bacterial features were categorised as ‘Definite Viral’ (DV) if a matching virus was identified, or otherwise as ‘Probable Viral’ (PV). Children without detected sterile-site bacteria and with inconclusive clinical features of viral or bacterial infection were classified as ‘Uncertains’ (U). We set a threshold of 60mg/L for the maximum CRP as a minimum for inclusion into the PB group, or a maximum for inclusion in the PV and DV groups. Inclusion in the DB group was irrespective of CRP. Patients failing the CRP threshold were categorised as ‘Uncertain’ (U), alongside the other patients in this group with inconclusive clinical features. Controls (C) had no current or recent (previous two weeks) infectious symptoms or immunisations, and no identified or probable chronic infectious or inflammatory conditions. CRP: C-reactive protein. WCC: white cell count

### Biomarker measurement

Heparinised blood samples were kept at 4C pending plasma recovery by centrifugation and storage at −80C. LCN2 and MMP-8 were measured using commercially available immunoassays (RD Systems, UK) following manufacturer’s instructions, by laboratory personnel blinded to the diagnostic group of each sample. Presence of visible haemolysis and the number of previous plasma freeze-thaw cycles prior to EIA had no systematic effect on measurement. The coefficient of variation for these measurements was 8.26% (calculated from 28 samples with 2 replicates). CRP and creatinine values contemporary with research blood sampling were obtained from hospital data for 156 children. CRP was measured by ELISA in controls (RD Systems, UK).

### Statistical analyses

Diagnostic performance of clinical markers was assessed using the area under the receiver operating characteristic (AUROC) curves to compare the sensitivity and specificity of selected markers. Cut-off values were chosen based on highest sensitivity and specificity to predict outcome (Supplementary Table 1). The diagnostic performance of biomarkers was assessed by logistic regression, using LCN2, CRP and MMP-8 as independent variables and the condition to diagnose as the dependent variable. A Chi-square statistic was used to test for equality of the AU-ROC (Stata 11, Stata Corporation) (6). Laboratory measures and ages in each patient group were compared using Kruskal-Wallis tests and Dunn’s pairwise comparisons. Frequency variables including sex, ethnicity and severity of illness were compared using the Fisher Exact or Chi-square test of independence, using Prism 5 (GraphPad Software Inc).

## Results

### Patient groups

Table 1 shows the demographic features of the patient groups. Ethnicity and gender did not differ significantly. Children in the DV group were younger than those in PB and C groups, as expected for viral respiratory infection. The proportion of children with severe illness was higher in the DB than DV group. *LCN2 and MMP-8 are associated with SBI.* Concentrations of plasma LCN2 and MMP-8 were lowest in controls, and increased stepwise with likelihood of SBI (Table 1). CRP, LCN2 and MMP-8 discriminated patients in the DB and DV or C groups (Figure 2a). We derived ROC curves for CRP, LCN2, MMP-8 for all 191 patients, and for neutrophil proportion and white cell counts in 175 patients with available values, based on the comparison of the DB group with the other febrile patients (combined PB, U and DV groups) (Figure 2b). LCN2 used as a continuous variable predicted confirmed SBI with an AUROC of 0.88 (95% CI: 0.82-0.94), matching the AUROC for CRP (0.89 [95% CI: 0.84-0.94]). MMP-8, neutrophil proportion and white cell count predicted confirmed SBI less well than LCN2 and CRP (AUROC 0.80, 0.71 and 0.69 respectively) (Figure 2b).

**Table 1.**
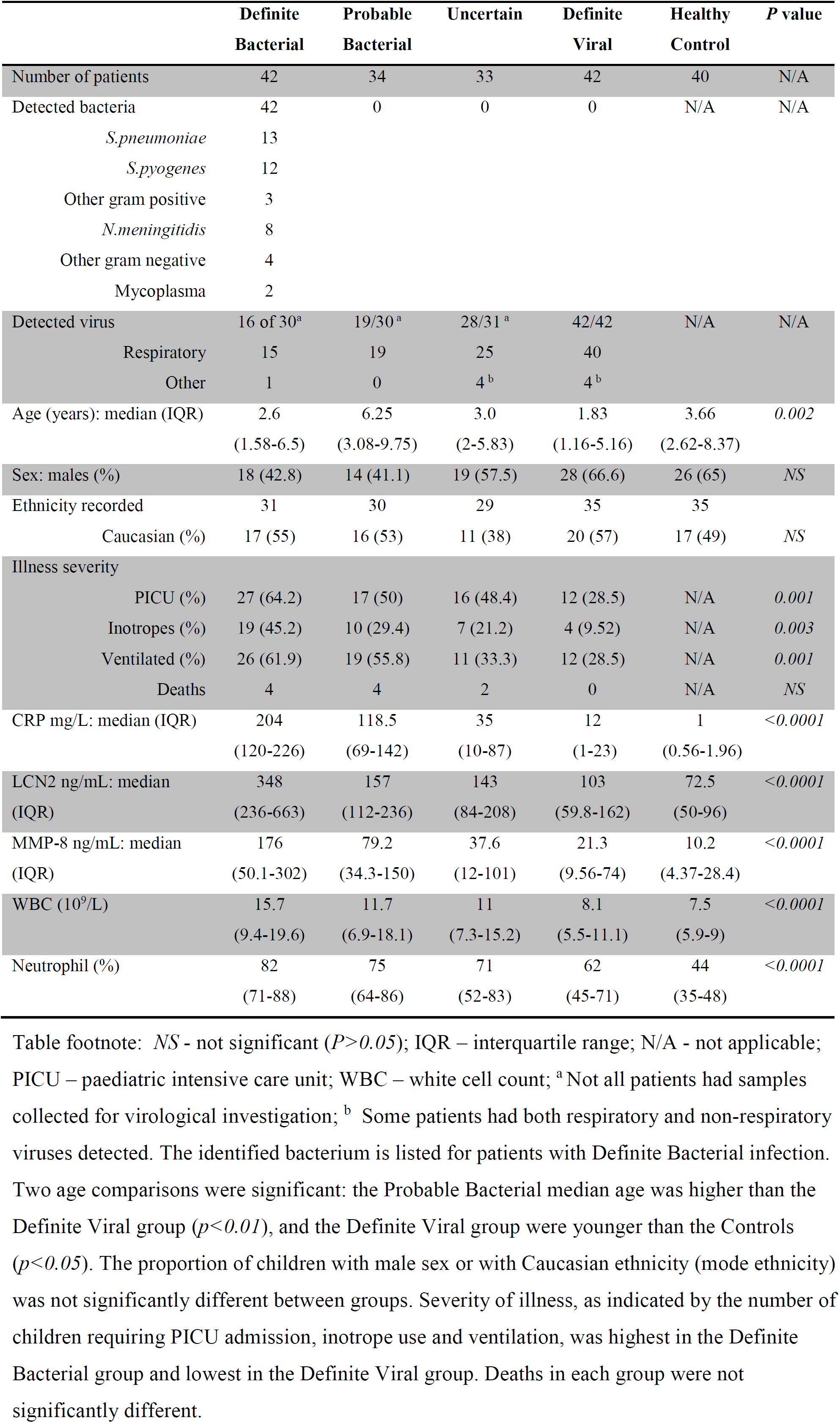
Demographic and clinical data of recruited subjects.

**Fig. 2.**
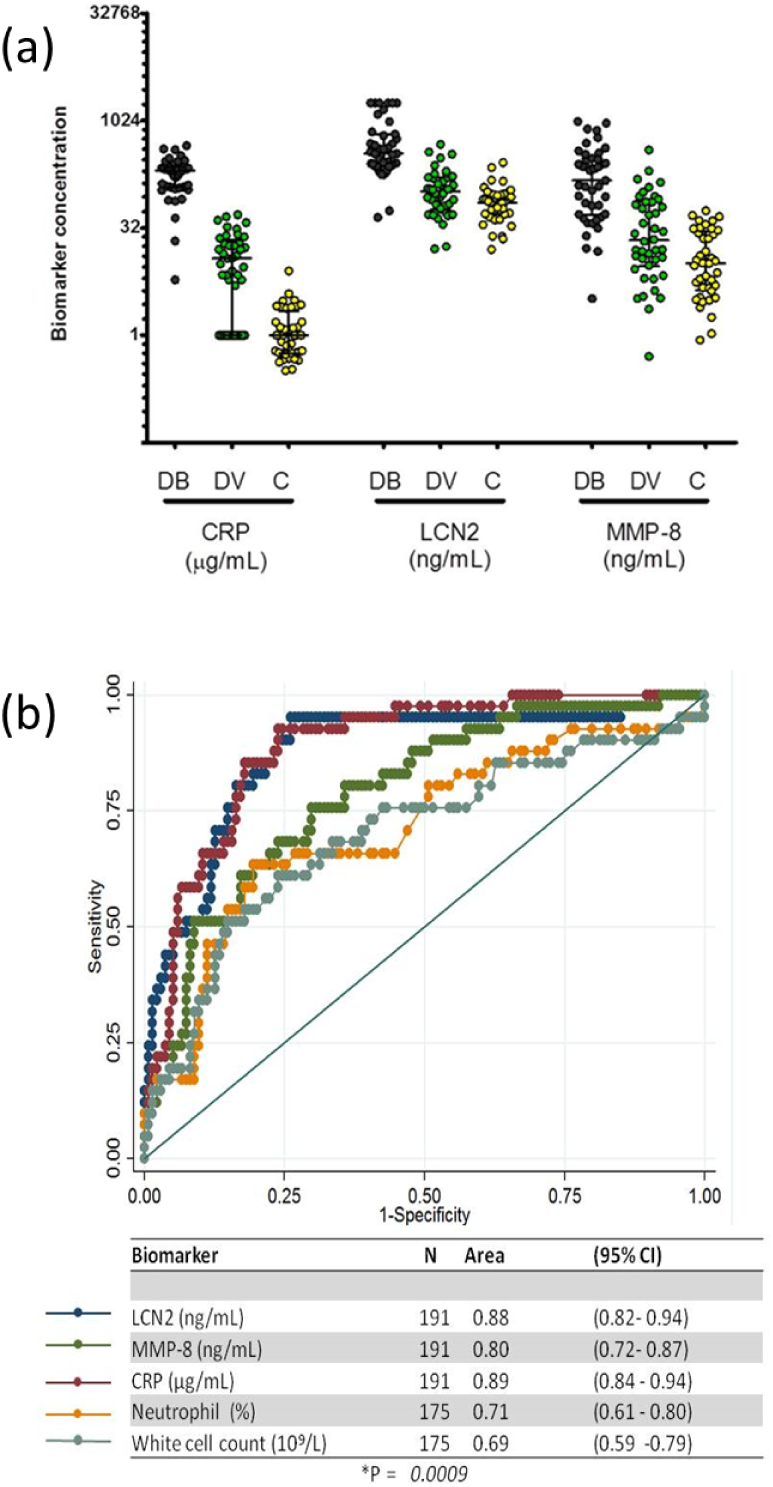
Distribution of LCN2 and MMP-8 concentration in the population studied. (A) Scatter plot of CRP, LCN2 and MMP-8 concentrations in patient groups DB, DV and C. (B) Diagnostic performance of LCN2, MMP-8 and CRP (calculated for 191 patients); and of neutrophil proportion and white cell count (calculated from 175 patients with available data) to predict SBI. Biomarker concentrations were used as continuous variables to predict confirmed bacterial infection and their diagnostic performance is shown in ROC plots as AUROC. Statistical significance for differences between the ROC curves was assessed on 175 patients with all variables were available (P=0.0009). DB – definite bacterial; DV – definite viral; C – healthy control; U – uncertain; PB – probable bacterial

### LCN2 performance is not confounded by creatinine concentration

As LCN2 levels increase with renal damage(7), we determined whether renal impairment influenced the diagnostic performance of LCN2. Creatinine levels contemporaneous with research bloods were available for 162 children (median and IQR: 39 and 34-50 µmol/L), and did not significantly differ between clinical groups (Kruskal-Wallis, P=0.36). The adjusted logistic regression analysis indicated that prediction of SBI by LCN2 was not affected by creatinine concentration (supplementary Table 1).

### Combination of LCN2 and CRP to predict SBI

CRP was used as a benchmark to evaluate the performance of LCN2 to predict confirmed SBI in 191 patients with available CRP and LCN2 data. The AUROC of LCN2 and CRP were strongly associated with confirmed SBI. Both biomarkers were highly correlated (r = 0.40, P<0.001). However, when both proteins were included into a logistic regression model, the combination was significantly superior to either marker alone (AU-ROC 0.92 [95%CI: 0.88-0.96]) (Figure 3 and Supplementary Table 1).

**Fig. 3.**
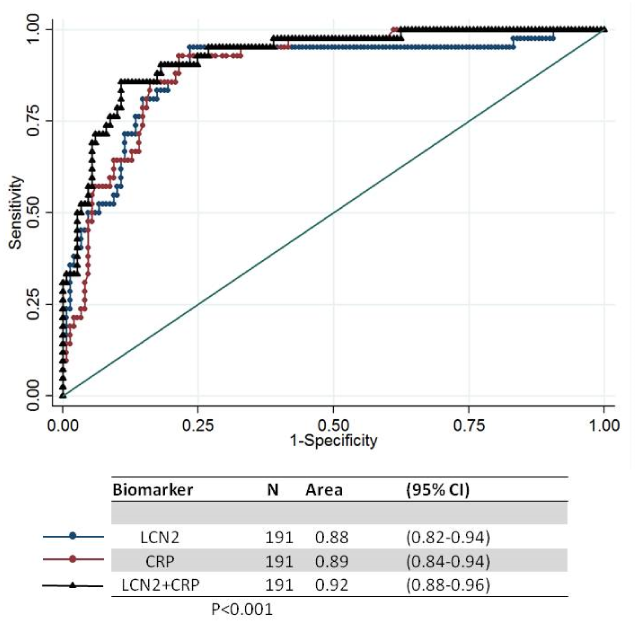
Diagnostic performance of CRP and LCN2 combined. The improved power of the combined biomarkers in regression analysis, and improved AUROC indicated that CRP and LCN2 each made an independent contribution to prediction of bacterial infection.

### LCN2 and CRP in uncertain cohorts

Without a sensitive gold-standard test for detection of SBI, the majority of SBI is ‘missed’, therefore we correlated LCN2 and CRP biomarker concentrations with clinical suspicion of SBI. There was a stepwise increase in biomarker concentration with increasing suspicion of SBI: from C, DV, U, PB, to B (Table 1, Figure 4).

**Fig. 4.**
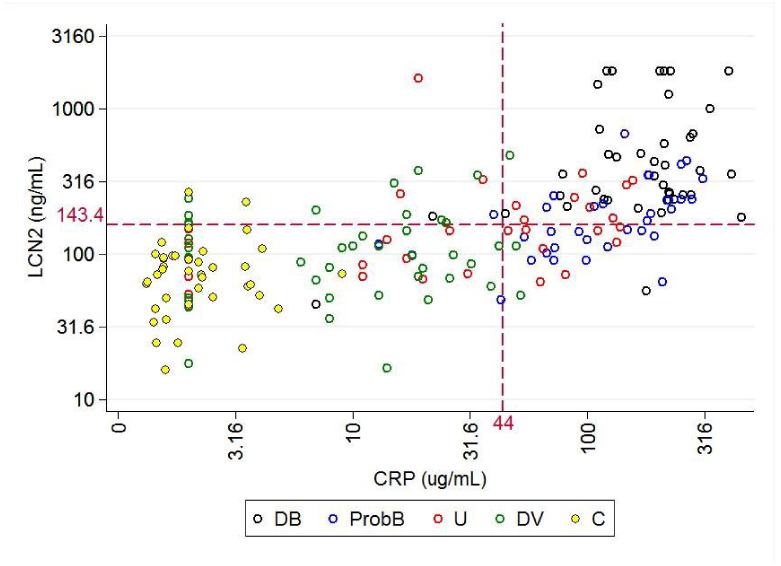
Scatter plot of LCN2 and CRP plasma concentration for all patients studied. Dashed lines indicate optimal cut-off values (highest sensitivity and specificity) for the combination of both biomarkers, which correspond to the 7th decile of the concentration of LCN2 and CRP (see Supplementary data).

We derived cut-off values for LCN2 and CRP based on the concentration that showed the highest sensitivity and specificity to discriminate confirmed bacterial infection (DB) from other febrile patients (Supplementary Table 1). The sensitivity and specificity of LCN2 to predict DB were 83.3% and 82.5%. CRP had sensitivity and specificity of 85.7% and 83.8%. When the threshold value for LCN2 was applied to the other groups, there was a stepwise increase in prediction of SBI: 2 of 40 (5%) (C), 6 of 42 (14%) (DV), 9 of 33 (27%) (U), 15 of 34 (44%) (PB). The sensitivity and specificity of both markers combined (equal or greater than their 7th decile) was 90.4% and 86.5%.

## Discussion

This study shows that LCN2 is a sensitive and specific biomarker associated with SBI in children with febrile illness and has potential to guide antibiotic treatment decisions. The best diagnostic performance was achieved by combining LCN2 and CRP (AUROC of 0.92).

LCN2 is a 21-kD glycoprotein secreted by neutrophils, hepatocytes and renal tubular cells and it is usually found at low concentration (20 ng/mL) in biological fluids (8). The role of LCN2 in the innate defence against bacterial infection has been attributed to its ability to interfere with bacterial iron uptake through competition with the bacterial siderophore enterobactin (9, 10). The role of LCN2 as a potential marker of bacterial infection as described in previous studies (5, 11).

Similarly, LCN2 gene expression has been shown to discriminate bacterial from viral infection, with 22-fold increased expression in bacterial patients compared to controls (12). Unlike CRP, the diagnostic performance of LCN2 is unaffected by age: LCN2 gene expression is central to an immune-metabolic network implicated in neonatal bacterial infection (13).

In comparison with CRP, LCN2 has a much shorter half-life (<20 minutes) (14), rising more rapidly than CRP in a study in neonates(15)and responding more rapidly to successful antibiotic treatment (11). Although a fast turnover is a desirable quality for a biomarker to monitor infection and treatment, it is also a diagnostic limitation when patients have received prior antibiotic treatment at the time of diagnosis.

LCN2 has recently emerged as a biomarker of acute kidney injury (7). We investigated if the association of LCN2 with SBI was confounded by impaired renal function. Adjusting the analysis for creatinine concentration did not affect the diagnostic performance of LCN2 to predict SBI, which suggests that neutrophils are the likely source of LCN2. The diagnostic performance of LCN2 exceeded that of MMP-8, previously proposed as a paediatric sepsis biomarker (16). In this study we have defined patient groups by a gradient from low-to high likelihood of SBI. This has the advantage to capture the real world situation in which the majority of patients have no definitively confirmed or excluded infection. The clinical group most likely to have SBI was defined by a positive blood culture (gold standard) and we used this variable as our predicted outcome to evaluate the diagnostic performance of LCN2 and MMP-8. Thus, it was not possible to compare diagnostic value of LCN2 to the gold standard; instead, we used CRP for comparison as the measurement of this biomarker to guide antibiotic treatment is commonly used in secondary and tertiary care.

We have shown that the biomarker concentration of LCN2 and MMP-8 was proportional to the estimated likelihood of SBI, with unconfirmed patients tending to have intermediate levels (47% of PB group had LCN2 > 175.3 ng/mL threshold). We did not compare the levels of LCN2 or MMP-8 to CRP in the groups with diagnostic uncertainty, as this comparison would be confounded by the use of the CRP result to allocate the patients to clinical groups (PB, U, DV). Procalcitonin is a promising biomarker of bacterial infection with kinetics comparable to those of LCN2 and it shown a modest advantage over CRP in the prediction of SBI (17), but we did not have these data for comparison.

Unless a biomarker is identified which discriminates SBI and non-SBI patients at non-overlapping concentrations, children with intermediate, ‘grey’ clinical markers of bacterial infection will remain as the group in whom the biomarkers are least effective, but most needed. The potential for LCN2 alone or in combination with CRP to improve the rapid identification of children with SBI at an early time, before culture results are available, could benefit clinical decision-making for children presenting with febrile illness, and improve antibiotic coverage in the easily missed minority of children with an SBI. The potential for LCN2 to improve antibiotic management decisions in febrile children should be evaluated in clinical trials.

## AUTHOR CONTRIBUTIONS

These contributions follow the Contributor Roles Conceptualization: JH, ML, CCP; Data curation: JH, HH, MLT, BK, CCP; Formal analysis: JH, ML, CCP; Funding acquisition: JH, ML, BK, CCP; Investigation: JH, VJ, MC, SG, MSH CCP; Methodology: JH, VJ, MC, SG, MSH CCP; Supervision: JH, ML, CCP; Writing – review & editing: all authors.

## COMPETING FINANCIAL INTERESTS

The authors declare no competing financial interests.

This work was supported by the Imperial College Comprehensive Biomedical Research Centre [DMPED P26077 to J.A.H.]; the Wellcome Trust Centre for Respiratory Infection at Imperial College; and the Southampton NIHR Wellcome Trust Clinical Research Facility and NIHR Wessex Local Clinical Research Network. CC-P is supported by the Medical Research Council UK (Clinician Scientist Fellowship: G0701885) and the NIHR (BRC/RCF: AC12/108).

We thank the patients and their families for taking part, Prof Saul Faust, Dr Sanjay Patel and Jenni McCorkell at the University of Southampton and Southampton University Hospital NHS Foundation Trust, and the clinicians and research teams for their support.

## Supplementary Material

**Supplementary Table 1.**
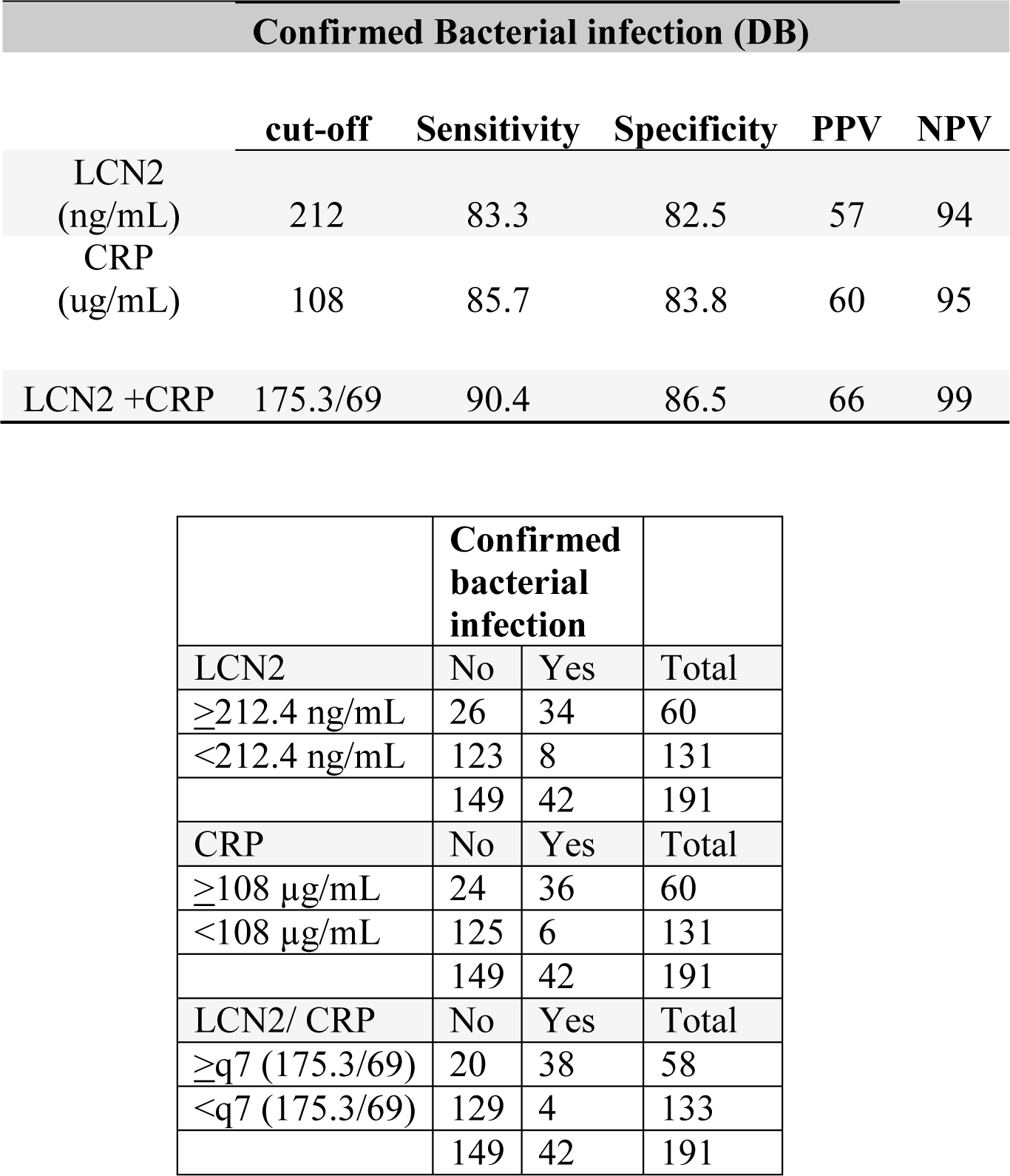
sensitivity and specificity of LCN2, CRP and combined LCN2 and CRP.

| **Confirmed Bacterial infection (DB)** |  |  |  |  |  |
| --- | --- | --- | --- | --- | --- |
|  | **cut-off** | **Sensitivity** | **Specificity** | **PPV** | **NPV** |
| LCN2<br>(ng/mL) | 212 | 83.3 | 82.5 | 57 | 94 |
| CRP<br>(ug/mL) | 108 | 85.7 | 83.8 | 60 | 95 |
| LCN2 +CRP | 175.3/69 | 90.4 | 86.5 | 66 | 99 |

|  | **Confirmed bacterial infection** |  |  |
| --- | --- | --- | --- |
|  | No | Yes | Total |
| LCN2 |  |  |  |
| ≥212.4 ng/mL | 26 | 34 | 60 |
| <212.4 ng/mL | 123 | 8 | 131 |
|  | 149 | 42 | 191 |
| CRP |  |  |  |
| ≥108 µg/mL | 24 | 36 | 60 |
| <108 µg/mL | 125 | 6 | 131 |
|  | 149 | 42 | 191 |
| LCN2/ CRP |  |  |  |
| ≥q7 (175.3/69) | 20 | 38 | 58 |
| <q7 (175.3/69) | 129 | 4 | 133 |
|  | 149 | 42 | 191 |

PPV positive predictive value, NPV: negative predictive value

Identification of the cut-off value with the highest balance of sensitivity and specificity to distinguish between definite bacterial infection (DB vs. non-DB) for LCN2, CRP and combined LCN2 and CRP. The combination of LCN2 and CRP had the highest sensitivity and specificity corresponded to values above the 7^th^ decile of LCN2 and CRP concentration.

**Supplementary Table 2.** Association of LCN2 with definite bacterial infection: logistic regression model adjusted for serum creatinine concentration. Serum creatinine was available for 161 patients.

| Outcome |  |  |
| --- | --- | --- |
| **Confirmed bacterial infection** |  |  |
|  | OR (95%CI)<br>[P value]<br>N of patients |  |
| Univariate logistic regression | crude | adjusted* |
| LCN2 (ng/mL) | 1.006 (1.003-1.009)<br>[<0.001]<br>N=191 | 1.005 (1.002-1.008)<br>[<0.001]<br>N=161 |

As LCN2 levels increase with renal damage [1], we determined whether renal impairment influenced performance. Creatinine levels contemporaneous with research bloods were available for 162 children (median and IQR: 39 and 34-50 µmol/L), and did not significantly differ between clinical groups (Kruskal-Wallis, *P*=0.36). The adjusted logistic regression analysis indicated that prediction of DB by LCN2 was not affected by creatinine concentration.

